# Genomic and proteomic analysis of Human herpesvirus 6 reveals distinct clustering of acute *versus* inherited forms and reannotation of reference strain

**DOI:** 10.1101/181248

**Authors:** Alexander L. Greninger, Giselle M. Knudsen, Pavitra Roychoudhury, Derek J. Hanson, Ruth Hall Sedlak, Hong Xie, Jon Guan, Thuy Nguyen, Vikas Peddu, Michael Boeckh, Meei-Li Huang, Linda Cook, Daniel P. Depledge, Danielle M. Zerr, David M. Koelle, Soren Gantt, Tetsushi Yoshikawa, Mary Caserta, Joshua A. Hill, Keith R. Jerome

## Abstract

Human herpesvirus-6A and -6B (HHV-6) are betaherpesviruses that reach >90% seroprevalence in the adult population. Unique among human herpesviruses, HHV-6 can integrate into the subtelomeric regions of human chromosomes; when this occurs in germ line cells it causes a condition called inherited chromosomally integrated HHV-6 (iciHHV-6). To date, only two complete genomes are available for HHV-6B. Using a custom capture panel for HHV-6B, we report near-complete genomes from 61 isolates of HHV-6B from active infections (20 from Japan, 35 from New York state, and 6 from Uganda), and 64 strains of iciHHV-6B (mostly from North America). We also report partial genome sequences from 10 strains of iciHHV-6A. Although the overall sequence diversity of HHV-6 is limited relative to other human herpesviruses, our sequencing identified geographical clustering of HHV-6B sequences from active infections, as well as evidence of recombination among HHV-6B strains. One strain of active HHV-6B was more divergent than any other HHV-6B previously sequenced. In contrast to the active infections, sequences from iciHHV-6 cases showed reduced sequence diversity. Strikingly, multiple iciHHV-6B sequences from unrelated individuals were found to be completely identical, consistent with a founder effect. However, several iciHHV-6B strains intermingled with strains from active pediatric infection, consistent with the hypothesis that intermittent de novo integration into host germline cells can occur during active infection Comparative genomic analysis of the newly sequenced strains revealed numerous instances where conflicting annotations between the two existing reference genomes could be resolved. Combining these findings with transcriptome sequencing and shotgun proteomics, we reannotated the HHV-6B genome and found multiple instances of novel splicing and genes that hitherto had gone unannotated. The results presented here constitute a significant genomic resource for future studies on the detection, diversity, and control of HHV-6.

**Author Summary:** HHV-6 is a ubiquitous large DNA virus that is the most common cause of febrile seizures and reactivates in allogeneic stem cell patients. It also has the unique ability among human herpesviruses to be integrated into the genome of every cell via integration in the germ line, a condition called inherited chromosomally integrated (ici)HHV-6, which affects approximately 1% of the population. To date, very little is known about the comparative genomics of HHV-6. We sequenced 61 isolates of HHV-6B from active infections, 64 strains of iciHHV-6B, and 10 strains of iciHHV-6A. We found geographic clustering of HHV-6B strains from active infections. In contrast, iciHHV-6B had reduced sequence diversity, with many identical sequences of iciHHV-6 found in individuals not known to share recent common ancestry, consistent with a founder effect from a remote common ancestor with iciHHV-6. We also combined our genomic analysis with transcriptome sequencing and shotgun proteomics to correct previous misannotations of the HHV-6 genome.

## Introduction

HHV-6 is a ubiquitous betaherpesvirus that is divided into two species (HHV-6A and -6B) (1). HHV-6B infects >90% of children by 2 years of age, causing roseola, also called exanthem subitem or sixth disease, which is the leading cause of febrile seizures among children (2–5). The virus persists in multiple cell types with consistent detectable viral DNA in saliva. HHV-6B reactivates in approximately 50% of allogeneic hematopoietic cell transplant (HCT) patients and is the most common cause of encephalitis in this setting. HHV-6B has also been associated with graft-versus-host disease, hepatitis, pneumonitis, and mortality after HCT, although causality remains to be proven (2).

Like other human herpesviruses, HHV-6A and -6B establish lifelong latency, but unique among human herpesviruses, they have the ability to integrate into human chromosomes. When this integration occurs in a germ cell, the virus can be passed to offspring and results in inherited chromosomally integrated HHV6 (iciHHV6). Affected individuals have a copy of the virus in each of their cells and the ability to pass on the integrated state to 50% of their offspring. IciHHV-6 is present in 0.5-2% of the population, constituting almost 70 million people worldwide, with the majority of these being iciHHV-6B (6). iciHHV-6 can also be passed between individuals via transplantation (7, 8). IciHHV-6 was recently associated with an increased risk of acute graft versus host disease and CMV viremia in HCT patients (9). Integrated virus has been shown to reactivate both *in vitro* and *in vivo* and can confound assays for active HHV-6 infection (10). The mechanism of integration and viral proteins required for integration are unclear (11).

Clinical testing for HHV-6 has hitherto been reserved to large academic medical centers and reference labs due to concerns over reactivation in HCT patients. Unbiased metagenomic sequencing has uncovered HHV-6 infection in a number of cases of encephalitis and febrile illness that were previously “unsolved” (12–14). Of note, HHV-6 has recently been included in new, rapid, point-of-care multiplex PCR panels for meningitis/encephalitis and febrile illness (15). Given the ease of use and extraordinarily rapid turn-around time of these multiplex PCR panels, they have already been adopted by thousands of hospitals across the world (15–17). Because it is not uncommon for children to have HHV-6B in their CSF around the time of primary infection, we expect the coming years to see hundreds of thousands of HHV-6 infections detected that previously would have gone undetected based on the sheer number of samples that will be tested for HHV-6 (18). With so many new infections detected, there is an increasing need to understand the clinical associations, sequence diversity, and basic biology of this virus

To date, only two complete genomes are available – the Z29 type strain from Zaire and the HST strain from Japan -- and limited comparative genomics studies have been conducted for HHV-6B (19–21). These two genomes have multiple conflicting annotations for gene and protein products. In addition, the annotated gene functions are mostly based on homology from cytomegalovirus, another human betaherpesvirus. Gene boundaries, protein sequences, and diversity of strains across time, place, and iciHHV-6 status are relatively unknown. These factors are critical for being able to perform molecular mechanism studies of viral pathogenesis (22).

Given that so little is known about HHV-6 genome diversity, gene/protein annotation, and gene/protein function despite its clinical disease associations, there is an opportunity to use agnostic technologies to rapidly annotate the HHV-6 genome. Large scale genome sequencing, RNA-Seq and ribosome profiling have previously been conducted in other human herpesviruses to discover new genes and proteins and to ascribe novel functions to known genes of these obligate intracellular parasites (23–27). Here we report the results of the first large-scale genome sequencing effort for HHV-6B with 125 near complete genomes along with reannotation of the genome with comparative genomics, transcriptomics, and proteomics. The results reveal limited sequence diversity among HHV-6B sequences with geographical clustering of HHV-6B sequences from acute infections and identical iciHHV-6B sequences among individuals without known recent common ancestry. RNA sequencing and shotgun proteomics combined with comparative genomic analysis enabled a consensus re-annotation of HHV-6B gene products that will serve as a resource for future clinical and basic science studies of HHV-6B.

## Results

### Global genomic diversity of HHV-6

In order to understand the genomic diversity of HHV-6, we performed capture sequencing of 125 strains of HHV-6B, comprised of 20 viral isolates from Japan, 35 isolates from New York, 6 strains from Uganda, and 74 strains of iciHHV-6 (64 species B, 10 species A) from HCT recipients or donors in Seattle (Figure 1A, Table 1). The HHV-6B oligonucleotides designed for capture sequencing could retrieve >99% of the HHV-6B genome, with less than 1% unresolved due to repetitive elements. The same panel was able to retrieve approximately 80% of the HHV-6A genome, again due to repetitive elements and in this case the reduced sequence identity with the HHV-6B oligonucleotide set (Figure 1B). Across the HHV-6B strains, the recoverable contiguous HHV-6B unique long (UL) region measured 119.6 kb, the N-terminal U86 contig measured 3.1 kb, the U90/91 contig measured 6.0 kb, and the U94-U100 contig measured 10.2 kb. The 10 HHV-6A strains assembled ranged from lengths of 60kb to 119kb with a median length of 118kb.

**Figure 1.**
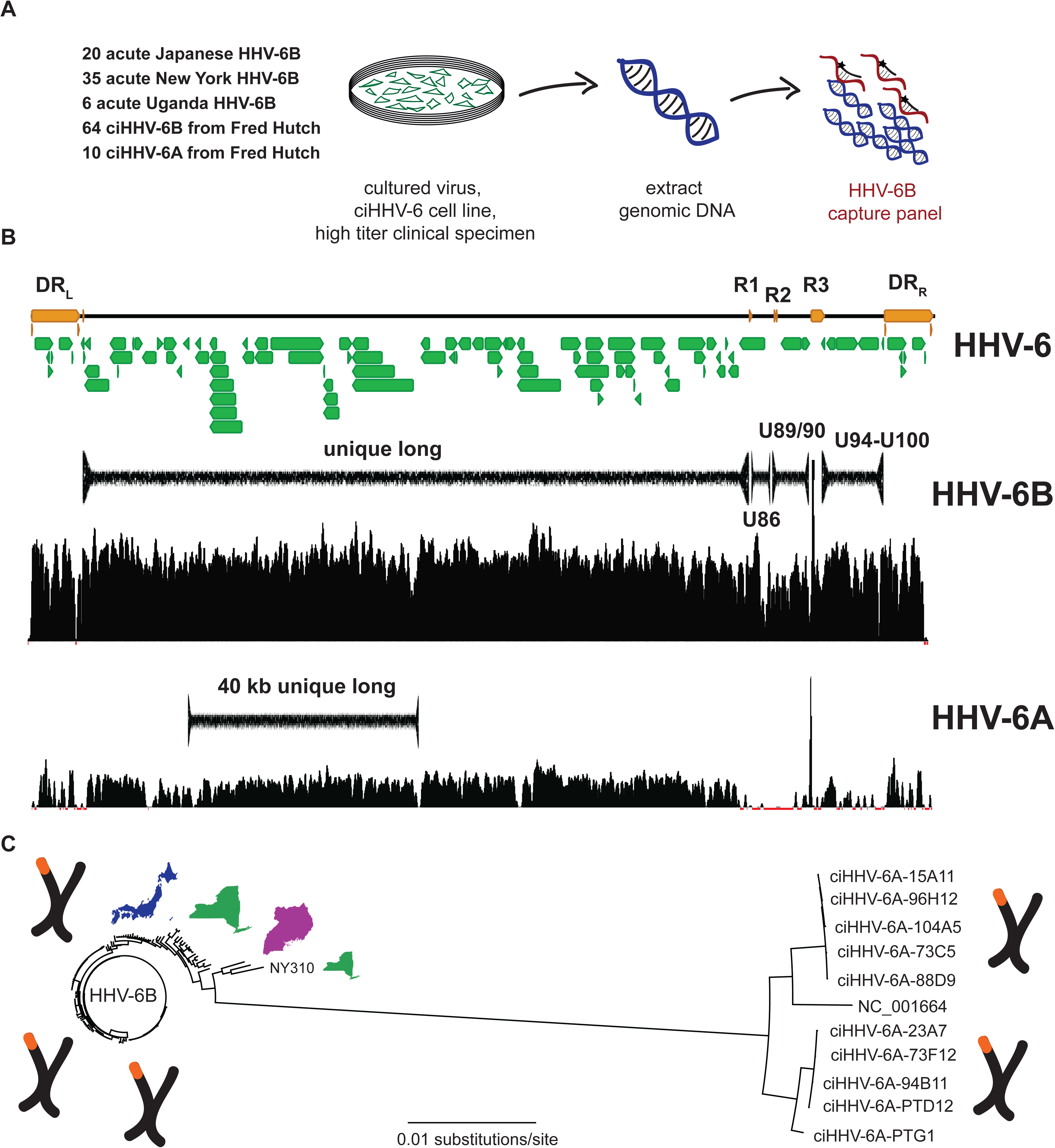
Experimental set up and HHV-6 genome calling mock up. A) A total of 129 HHV-6 specimens comprised of 55 cultured HHV-6B strains from acute infections, 6 clinical samples from acute infections, and 64 iciHHV-6B and 10 iciHHV-6A cell lines were sequenced using a capture panel based on the HHV-6B reference genome (NC_000898). B) Consensus genomes used for phylogenetic analysis were called for regions outside of the repeat regions for the HHV-6B specimens, including the unique long region (119kb), U90/91 region (6kb, between R2 and R3 repeats), and U94-100 (10kb, between R3 and DR-R repeat). The HHV-6B capture panel recovered much of the unique long region from HHV-6A specimens. C) Overall phylogeny of 40.2kb sequence that was recovered from HHV-6A and HHV-6B strains sequenced reveals separation of HHV-6A and HHV-6B as separate herpesvirus species.

**Table 1.** Summary of Samples Sequenced in This Study

| Location | Number | Species | Clinical | Median Age |
| --- | --- | --- | --- | --- |
| Japan | 10 | HHV-6B | Acute, BMT | 28 yrs [3 - 64 yrs] |
| Japan | 10 | HHV-6B | Acute, Exanthem subitum | 1 yr [8 mo - 3 yrs] |
| New York | 35 | HHV-6B | Acute, fever | 11 mo [1-25 mo] |
| Fred Hutch | 64 | HHV-6B | ciHHV-6, BMT | 40 yrs [1 - 68 yrs] |
| Fred Hutch | 10 | HHV-6A | ciHHV-6, BMT | 57 yrs [21 - 63 yrs] |
| UW Virology | 11 | HHV-6B | reactivation, acute | unknown |
| Uganda | 6 | HHV-6B | Primary | unknown |

Abbreviations: BMT, bone marrow transplant

### Demographic characteristics of cohorts

The median age of the roseola cohort from Japan was 12 months [8 - 24 months], the New York febrile infant cohort was 11 months [1 – 25 months], and the Uganda cohort was 25 months. All 20 patients from the two cohorts from Japan were of Japanese ancestry and all 6 patients from the Uganda cohort were Black Africans. In the New York cohort 16/35 (45.7%) of patients were Caucasian, 8/35 (22.9%) were African-American, 4/35 (11.4) were Hispanic, 1/35 (2.9%) were Hispanic, and 6/35 (17.1%) were of unknown ethnicity. Of the iciHHV-6 samples sequenced, 68/74 (91.9%) of patients came from the United States, while 2 patients came from the United Kingdom, 2 patients from Germany, and 1 from Australia (S1 Table). The median age of iciHHV-6B individuals sequenced was 40 years [1 – 68 years] and 57 years [21 – 63 years] for iciHHV-6A individuals (Table 1).

### Comparison of HHV-6A and HHV-6B

Phylogenetic analysis of a 40.2 kb segment ranging from U18 to U41 that could be captured in both the HHV-6A and HHV-6B strains sequenced in this study demonstrated separate clustering of the HHV-6A and HHV-6B strains, consistent with their designation as unique species of human herpesviruses (Figure 1C). Recombination analyses using all 10 iciHHV-6A partial sequences, HHV-6A type strain, and 14 selected HHV-6B sequences revealed no recombination sites between HHV-6A and HHV-6B sequences. Two individuals from Germany and the United States who shared no relations were found to have identical iciHHV-6A sequences. HHV-6A sequences showed little divergence in this 40.2 kb region with 98.4% of sites having no nucleotide variants.

### Sequence divergence in HHV-6B

Phylogenetic analysis of the unique long region revealed a cluster of two viruses from Uganda and one from New York NY310 that comprise the most divergent HHV-6B viruses sequenced to date (Figure 1C). NY310 most closely aligned to the Z29 strain, differing from the Z29 strain by 644 of 119,635 sites (0.54%). NY310 showed greater genetic distance to the next closest American strain NY434 (703 sites, 0.59%). This strain had no obvious unique demographic or clinical characteristics, having been derived from an 18-month old white male with fever after only 2 passages in culture (S1 Table). NY310 served as the outgroup for all subsequent phylogenetic analyses of HHV-6B genomes.

Overall, the 119.6kb HHV-6B unique long contig showed remarkably little sequence divergence with 98.1% of sites being identical among all 127 HHV-6B genomes and an overall pairwise identity of >99.9% between strains. The prototypical typing gene U90 was the most divergent with changes at 6.14% of total sites, while U15 and B6 repeat genes had the least amount of divergence with only 0.52% and 0.41% of sites being divergent (Figure 2). Average nucleotide diversity of the Ugandan HHV-6B UL sequences was 7 times that of the iciHHV-6B strains, which had the least amount of diversity of all strains profiled (Table 2). The Japanese isolates were approximately 40% less diverse than New York isolates. Both Achaz and Tajima neutrality tests to test for non-random sequence evolution across the entire UL region were strongly negative in all cohorts, likely due to the population structure and demographic history of the samples analyzed (28, 29).

**Figure 2.**
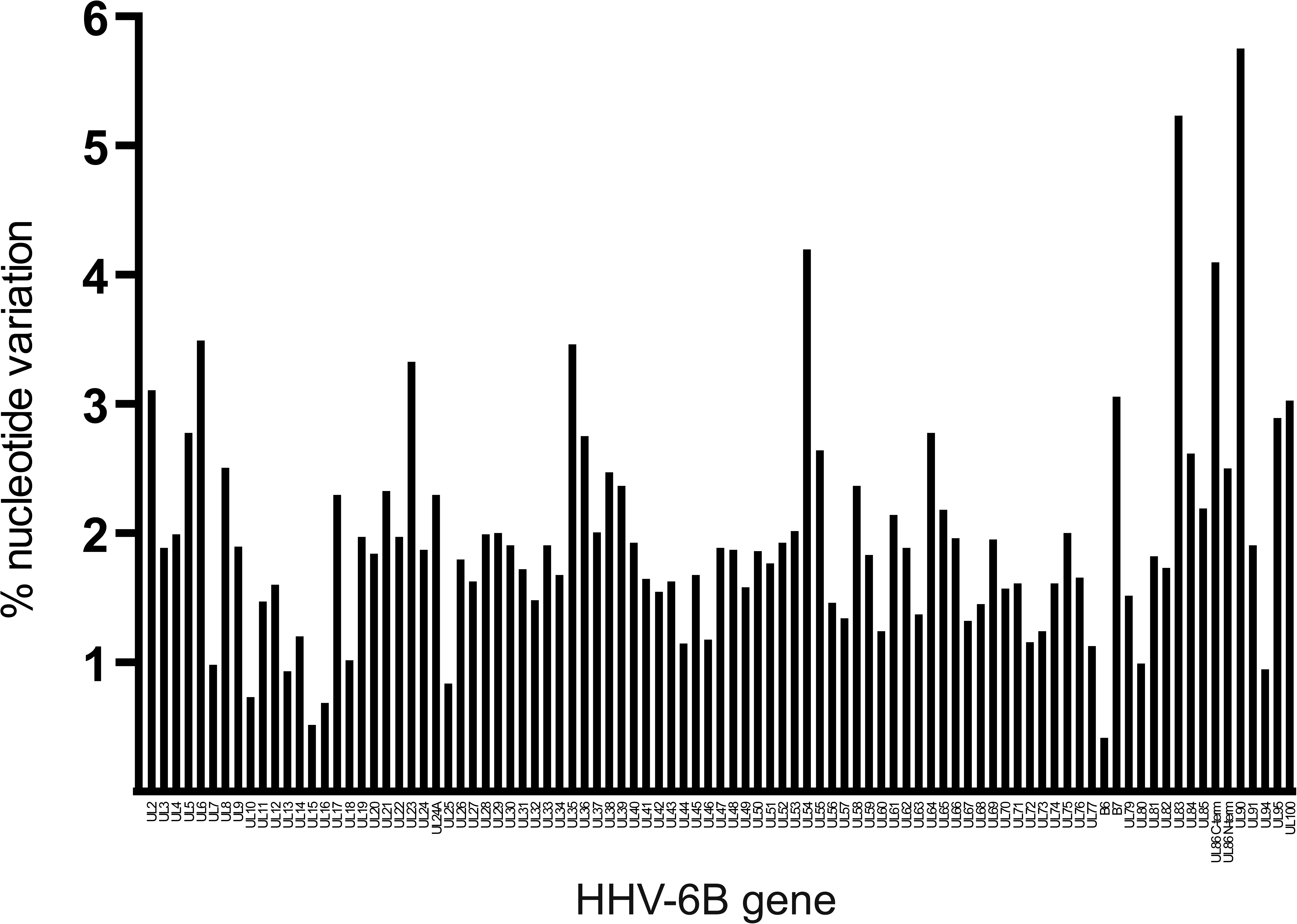
Nucleotide diversity by gene. Non-identical sites listed by gene by percent of sites with any variance. The prototypical typing gene UL90 is the most divergent with changes at 6.14% of total sites, while UL15 and B6 repeat genes had the least amount of divergence with only 0.52% and 0.41% of sites being divergent.

**Table 2.** Population genomics statistics for UL region by cohort sequenced

| Locus | Cohort | Samples | Nucleotide diversity | Achaz Y | Tajima's D | H-K sites |
| --- | --- | --- | --- | --- | --- | --- |
| 119kb UL | Japan | 20 | 109.5 | -0.75 | -0.85 | 25 |
|  | New York | 35 | 177.3 | -0.33 | -1.72 | 103 |
|  | Uganda | 6 | 357.3 | -0.39 | -0.11 | 34 |
|  | iciHHV-6B | 64 | 51.1 | -0.94 | -1.86 | 20 |
|  | All ULs | 127 | 144.9 | -1.59 | -2.02 | 162 |
| 40kb UL | 11 HHV-6A<br>129 HHV-6B | 140 | 256.0 | -1.59 | -1.47 | 144 |

### Phylogenetic analysis of HHV-6B sequences from acute infections

The continual replication of HHV-6B virus present in acute infections suggests a different evolutionary history than that of potentially preserved iciHHV-6B sequences, which are potentially preserved over time due to the high fidelity of human genomic replication. Phylogenies of the 119.6 kb unique long region of the HHV-6B genomes revealed clustering by geography as well as by sample type (i.e. acute HHV-6B infection versus iciHHV-6B) (Figure 3). Strains from active New York infections demonstrated the greatest amount of sequence divergence with at least two different clusters while Japanese strains all clustered together. Three of the Uganda sequences clustered among one of the New York strain clusters, while three others formed a unique clade (Figure 3). Sequence from the only known patient of Asian descent from New York (NY379) fell into the Japanese cluster along with two additional New York HHV-6B strains.

**Figure 3.**
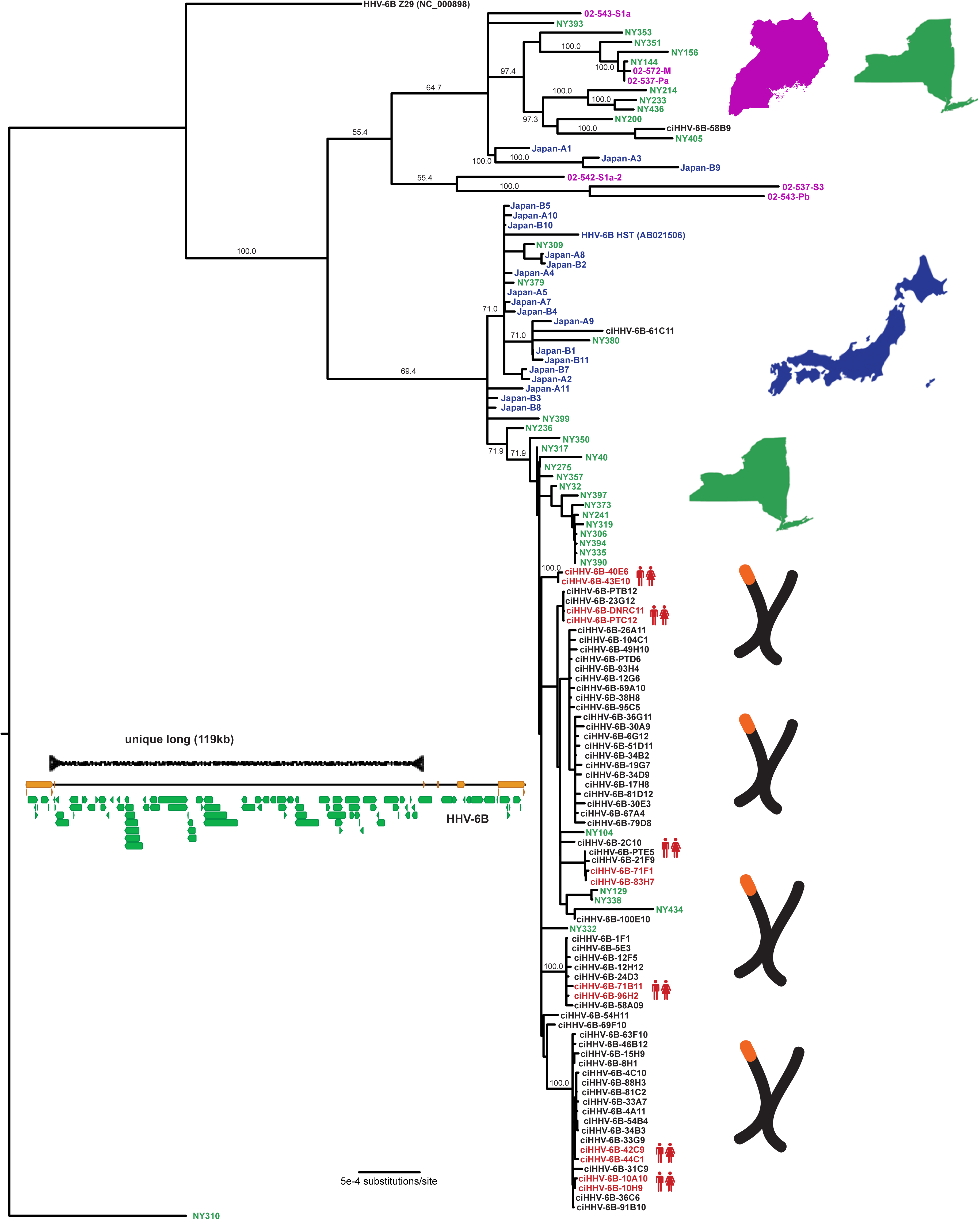
Phylogenetic tree of unique long region from HHV-6B samples. HHV-6B genomes were aligned using MAFFT, curated for sequence outside of repeat regions, and phylogenetic trees were constructed using MrBayes along the 119kb unique long region. HHV6-6B NY310 was used as an outgroup. Samples are colored and labeled for origin based on New York (green), Japan (blue), Uganda (purple), or iciHHV6 from HSCT recipients or their donors in Seattle (black), as well as whether two genomes were recovered from first-degree relatives (red).

### Identical iciHHV-6B strains in unrelated individuals

IciHHV-6B strains showed remarkable relatedness among unrelated individuals. Across 62 of the 64 iciHHV-6B UL regions sequenced here, only 334 of 119,635 (0.28%) sites had polymorphisms. Among iciHHV-6B HCT recipients whose donors were first-degree relatives, all 6 pairs had iciHHV-6B strains that were found to be identical (Figure 3). Identical iciHHV-6B strains were also found between unrelated individuals from Germany and the United States. Notably, resequencing of several of the identical iciHHV-6B strains from unrelated individuals gave identical sequence in 11 of 12 samples, controlling for possibility of laboratory contamination or sample mix-up (Figure S1). The lone outlier was a strain with a singular base with a variant allele frequency of almost exactly 50% in each sequencing replicate. Analysis of the off-target human mitochondrial reads from unrelated individuals revealed unique mitochondrial SNPs, confirming that these are from unrelated individuals (data not shown). IciHHV-6B strains were found intersperse among a New York cluster of acute infections as well as the Japanese cluster of acute infections, and several New York acute infection strains fell in the iciHHV-6B clusters. Branch lengths were generally longer for most of the acute infection strains indicating greater sequence divergence from common ancestor compared to iciHHV-6B strains.

### Sequence diversity of HHV-6B in non-UL regions, U90-91 and U94-100

Based on our data showing U90 to be the most divergent gene in HHV-6B, we sequenced an additional 11 U90 sequences from HHV-6 positive clinical specimens present in the UW Virology clinical lab. Phylogenies from the U90-91 and U94-100 regions revealed a similar topology to that of the unique long phylogeny with a few notable exceptions. New York strains again showed the greatest diversity and the Japanese strains again clustered together. The U90-91 phylogeny showed two Japanese strains (B1 and B4), a New York strain (NY40), a UW clinical isolate (UW_AH1), and one iciHHV-6B strain (61C11) that clustered with the Z29 type strain and four strains with U90 sequence present in Genbank (Figure 4, Figure S2B). Additional recent UW clinical strains for which U90 sequence was available clustered throughout the HHV-6B U90 tree recovered from the whole genome sequence, with one additional prominent outgroup (UW_BF2). The Japanese and New York strains were each located in their respective unique long cluster, while the iciHHV-6B-61C11 strain fell in the Japanese cluster. The disparity between the UL and U90 phylogenies is evidence of potential recombination in these strains with HHV-6B that is closer to the Zairian Z29 strain or the NY310 outgroup. Of note, the Japan-B1 strain also fell in a unique position in the U94-100 region among an iciHHV-6B cluster, while the NY40 strain was located in the U94-100 Japanese cluster (S2B Figure). In addition to the phylogenetic analysis indicative of recombination, Hudson-Kaplan RM estimates of parsimonious recombination events across the UL region ranged from 20 recombination sites for iciHHV-6B strains to 103 sites for New York strains (Table 2), suggesting widespread recombination within HHV-6 species.

**Figure 4.**
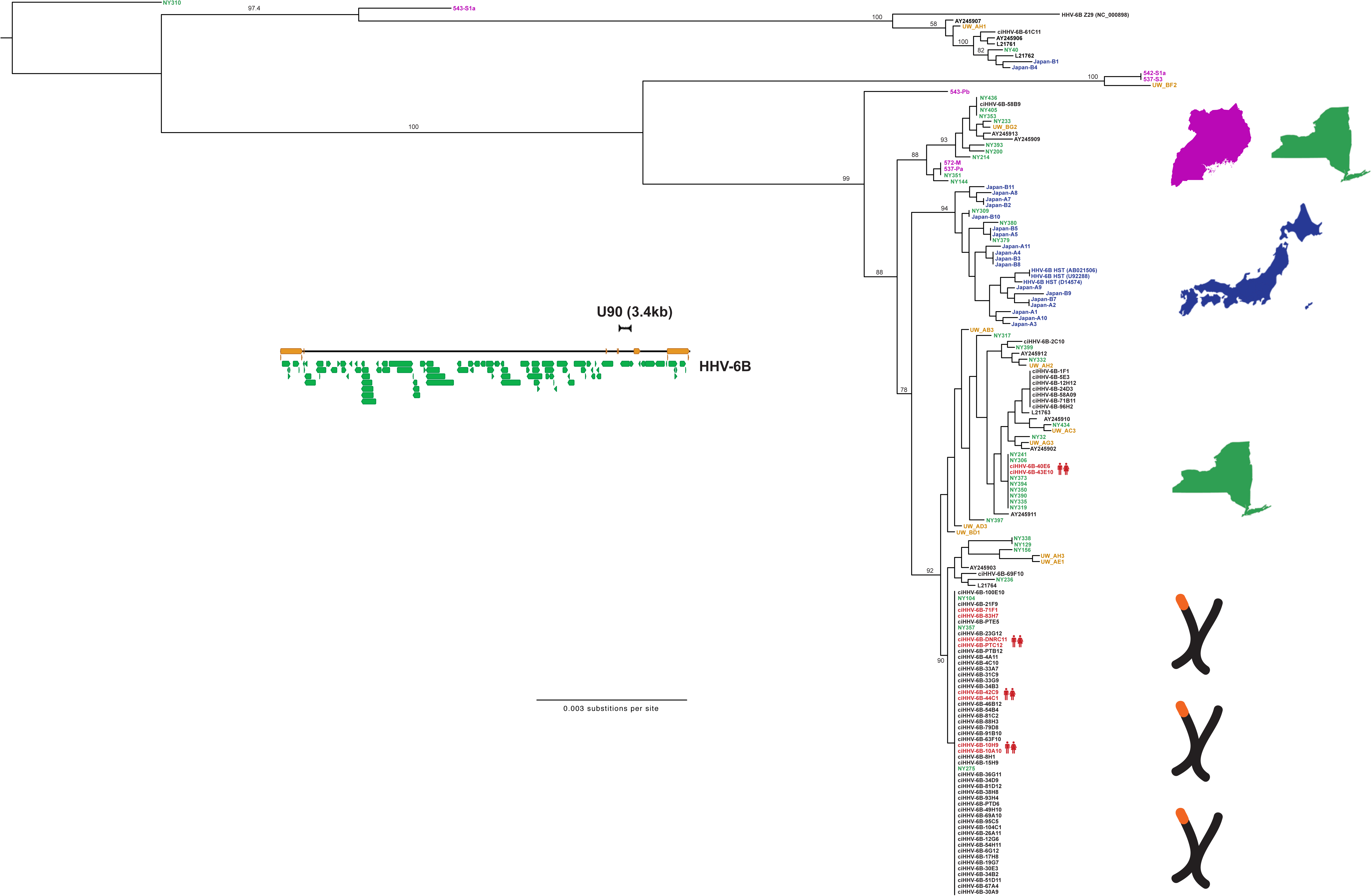
Phylogenetic trees of HHV-6B samples in U90 region including UW clinical isolates. HHV-6B U90 sequence captured from the 125 complete genomes and directly PCR-amplified from the UW cohort specimens were aligned using MAFFT and phylogenetic trees were constructed using MrBayes. Samples are colored and labeled for origin based on New York (green), Japan (blue), Uganda (purple), UW Virology clinical specimens (gold), or iciHHV6/FHCRC (black), as well as whether two genomes were recovered from first-degree relatives (red).

### Annotation of HHV-6B via comparative genomics

Multiple gene annotation discrepancies exist between the published HHV-6B Z29 and HST genomes. With the availability of 125 new HHV-6B genomes, we next examined the sites of these annotation discrepancies in our new HHV-6B genome sequences. For instance, the U91 gene contains an annotated splice site in the Z29 while no such annotation is found in the HST gene. Sanger sequencing of U91 cDNA from our lab’s cultured Z29 strain (Z29-1) revealed a different splice site 13 bp away from the annotated Z29 splice site, adding 5 additional amino acids to the middle of the U91 protein (Figure 5A). Both the cloned splice site and the annotated splice site contained canonical intronic splice sequencing (GU…AG). Cloning of the Z29-1 cDNA with the new splice site revealed an early stop codon that would disrupt the annotated C-terminal half of the protein in Z29 strains. Shotgun genomic sequencing of the cultured HHV-6B Z29-1 strain matched the Z29-1 cDNA sequence. Of note, Z29 is the only HHV-6B strain in our genomic sequencing with a single adenine insertion near the start of the second exon. Using the cloned splice site, all other U91 genes sequenced in this study would be in-frame to the end of the annotated U91, revealing that Z29 is likely unique among HHV-6B strains in missing the C-terminal half of U91.

**Figure 5.**
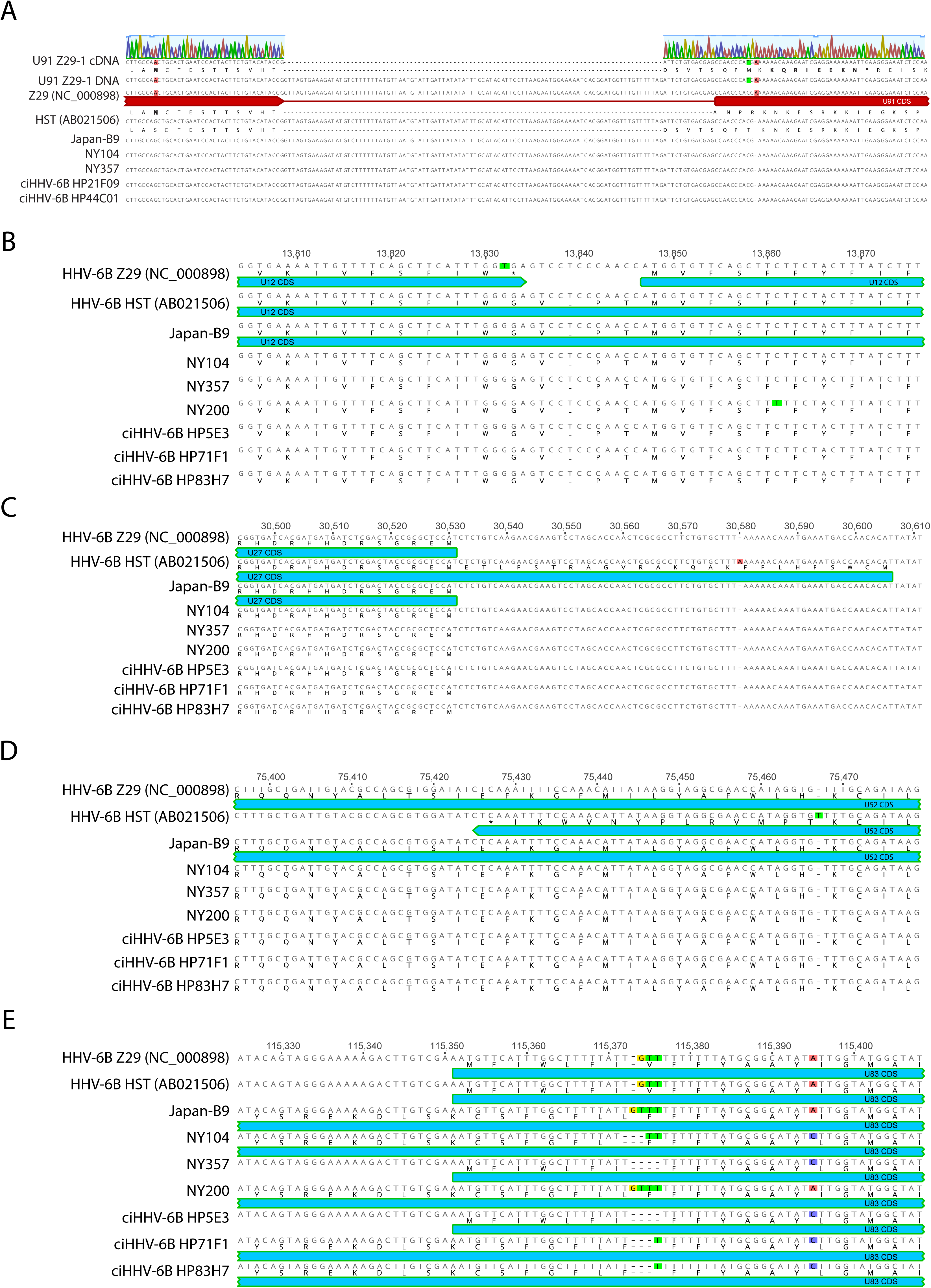
HHV-6B annotation based on comparative genomics. Differences in annotation between HHV-6B Z29 and HST sequences are compared with a subset of the 119 genomes sequenced in this study. A) Sanger sequencing of U91 cDNA revealed a different splice site 13-bp upstream than that which is annotated in the reference Z29 strain. The aberrant splice-site annotation in Z29 is likely due to a single base insertion found only in Z29 that alters the reading frame in the second exon. Genome sequencing of our cultured HHV-6B strain (Z29-1) confirmed the Z29-1 cDNA sequence. The reading frame depicted for Z29 is as annotated in the NCBI reference genome (NC_000898). Based on the newly discovered splice site, the Z29 U91 would contain an early stop codon while all other U91 sequences obtained in this study would continue the reading frame to the end of U91 as annotated in Z29. Several key different loci in U12 (B), U27 (C), and US52 genes (D) that alter the length of open reading frames in Z29 and HST are depicted. E) A homopolymeric polymorphism in U83 changes in expected length and sequence of its open reading frame between different strains.

Several other annotation discrepancies between exisiting HHV-6B sequences could be reconciled with our new genome sequences. The U12 gene in the Z29 strain is interrupted by a stop codon while the HST strain contains one long ORF (Figure 5B). Comparison with the 126 UL genomes sequences in this study show that for U12, the HST CDS should be considered the more representative of the original two genomes. Alternatively for U27 and U52, homopolymeric SNPs in HST creates abnormally long and short annotated ORFs, respectively, that are not reflected in the newly sequenced genomes (Figure 5C/D). Homopolymeric SNPs are also found in the U83 gene resulting in a polymorphic annotation across many of the sequence genomes (Figure 5E).

### Reannotation of HHV-6B genome through RNA-Sequencing and Shotgun Proteomics

Based on the number of discrepancies between HST and Z29 strain annotation that could be resolved by comparative genomics, we pursued RNA sequencing of the transcriptome of the HHV-6B Z29 type strain to more exhaustively reannotate the HHV-6B genome. Two biological replicates were prepared for the HHV-6B Z29 RNA-Seq library in SupT1 cells and were sequenced at coverages of 266X and 3600X, while one strand-specific library was prepared for a HHV-6B Z29 infected MOLT3 cells at an average coverage of 5751X. RPKM values for HHV-6 genes from SupT1 replicates were highly reproducible (r^2^ = 0.92) (Figure 6A). Compared to the Z29 transcriptome in SupT1 cells, the Z29 transcriptome in MOLT3 cells demonstrated significantly less correlation (r^2^ = 0.66) (Figure 6B). While only 3/104 (2.9%) HHV-6B CDS had 2-fold higher expression in in SupT1 cells compared to MOLT3 cells, 19/104 (18.2%) CDS had greater expression in MOLT3 cell lines (Figure 6C).

**Figure 6.**
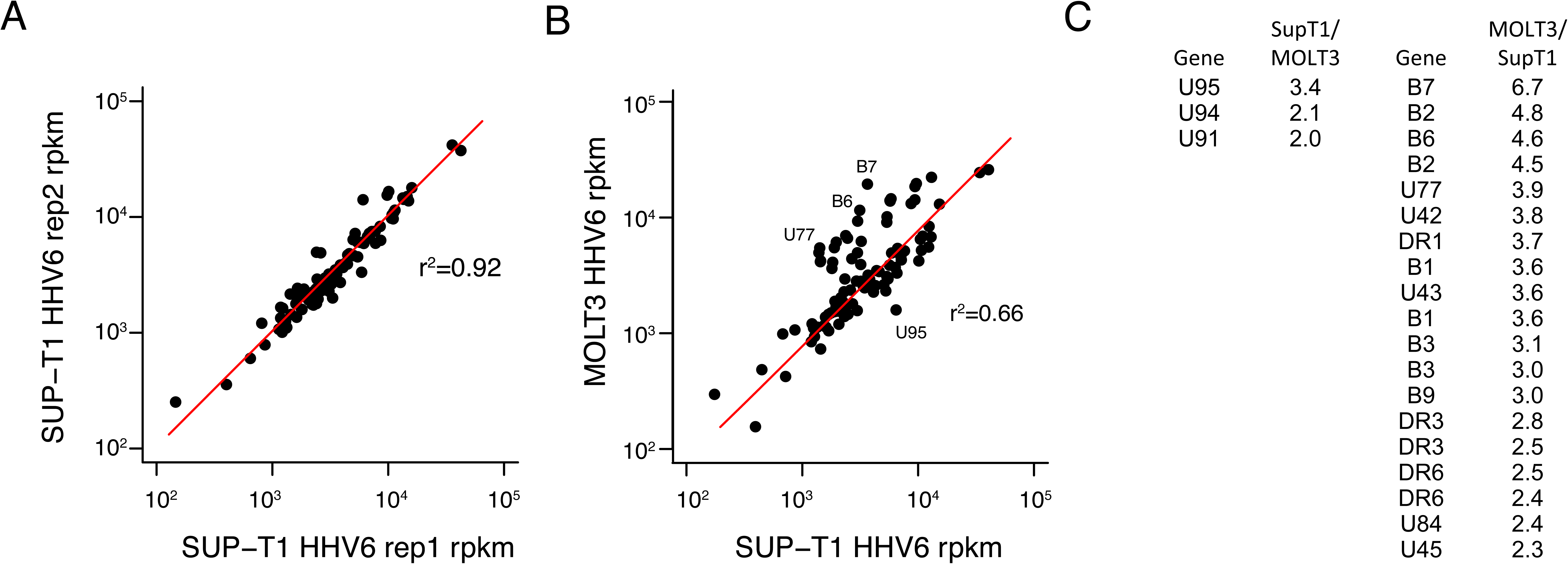
RNA Sequencing of Sup-T1 and MOLT3 cell lines asynchronously infected with HHV-6B Z29 type strain. RPKM values for HHV-6B Z29 transcripts in biological replicates of virus grown in Sup-T1 cells show excellent reproducibility (A). RPKM values of HHV-6B Z29 transcripts for virus grown in MOLT3 cells show differences in expression compared to virus grown in Sup-T1 cells (B). List of HHV-6B CDS with >2-fold absolute variation in expression in Sup-T1 and MOLT3 cell lines (C). Substantially more HHV-6B genes had higher expression in MOLT3 cells than in Sup-T1 cells.

Analysis of the mapped reads revealed a number of novel spliceoforms that were present. All splice sites mapped were perfectly conserved in the 127 HHV-6B genomes analyzed. Five of 43 (11.6%) total splice sites recovered were non-canonical with 4/5 (80%) non-canonical splice sites occurring in U7-U9 transcripts. To validate these novel spliceoforms and extensions that affected coding sequences, we performed shotgun mass spectometry on 1D gel-separated proteins from HHV-6B Z29 cultured in SupT1 cells (S3/4 Figure). Shotgun proteomic analysis produced 350 unique spectra covering 39 different HHV-6 proteins that may be viewed in MS Viewer (S2 and S3 Table).

Intriguingly, three novel U79 mRNA isoforms were found, one of which also demonstrated divergent splicing based on culture in SupT1 versus MOLT3 cell lines (Figure 7). Peptide confirmation of the novel U79 spliceoform present in SupT1 cells was confirmed with two peptides – LSTCEYLK with m/z 507.25 (2+), and YLCVR 355.68 (2+) – from shotgun proteomics analysis (S3 Table). The U19 gene demonstrated an unannotated splice junction just prior to the annotated stop codon, extending the C-terminus of the protein by 13 amino acids (Figure 8). Peptides immediately before and after the splice junction were recovered, confirming the expression of the C-terminal extension (DFLEEIAN 475.72 (2+) and SPENAVHESAAVLR 493.92 (3+) in S3 Table). Antisense reads along with a novel stop codon were recovered to the existing U83 annotation (Figure 9).

**Figure 7.**
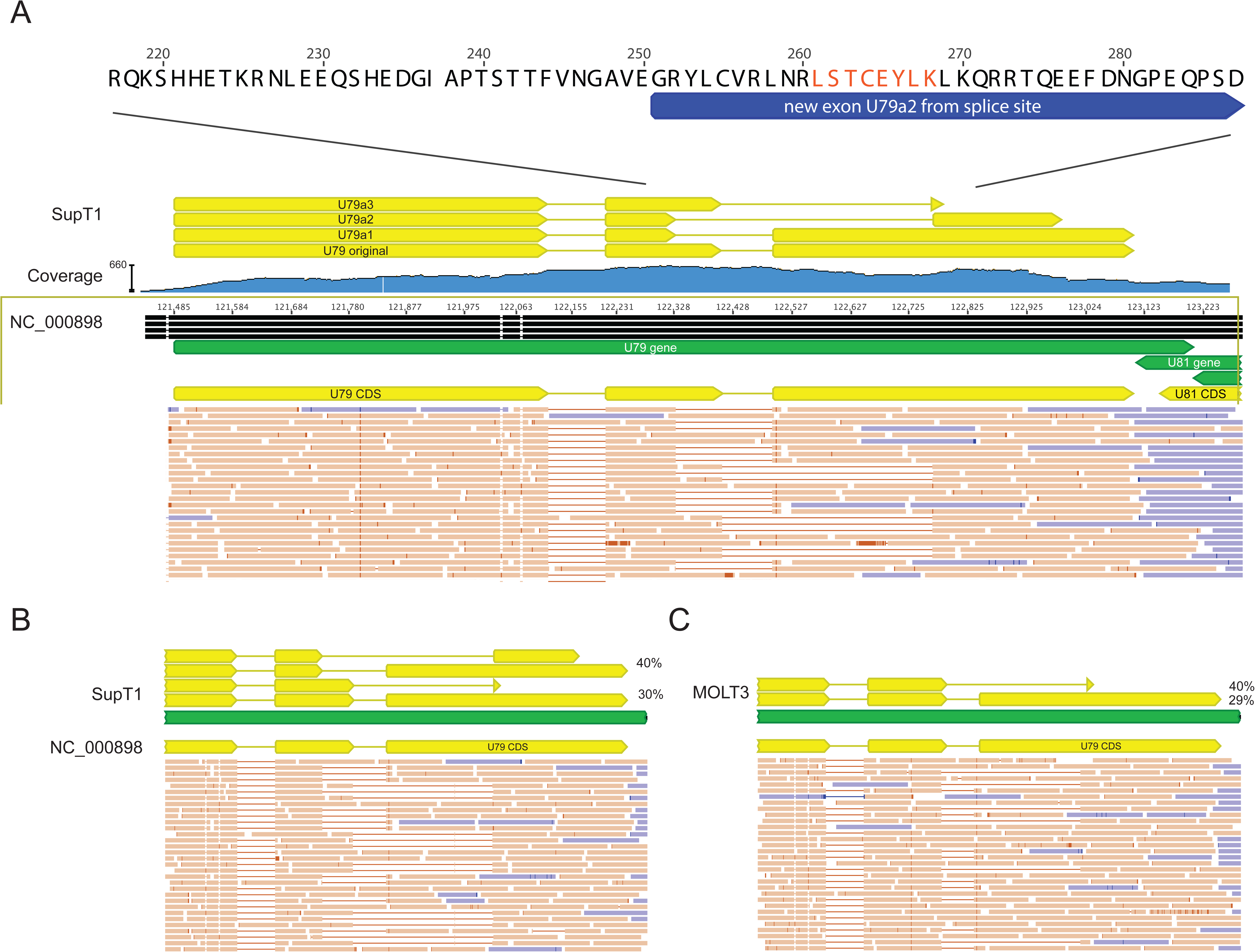
Alternative and differential splicing of HHV-6B U79 transcripts in Sup-T1 versus MOLT3 cells. Strand-specific RNA sequencing reveals three additional spliceoforms of the U79 gene in HHV-6B Z29 strain cultured in Sup-T1 cells compared with the Z29 reference annotation in NC_000898 (A). Reads depicted in orange are positive-sense reads, while negative-sense reads are shown in blue. The highlighted peptide from the U79a2 transcript in red was confirmed by shotgun proteomics of the Sup-T1 cultured HHV-6B. In MOLT3 cells, only two forms of splicing in U79 are detected (C).

**Figure 8.**
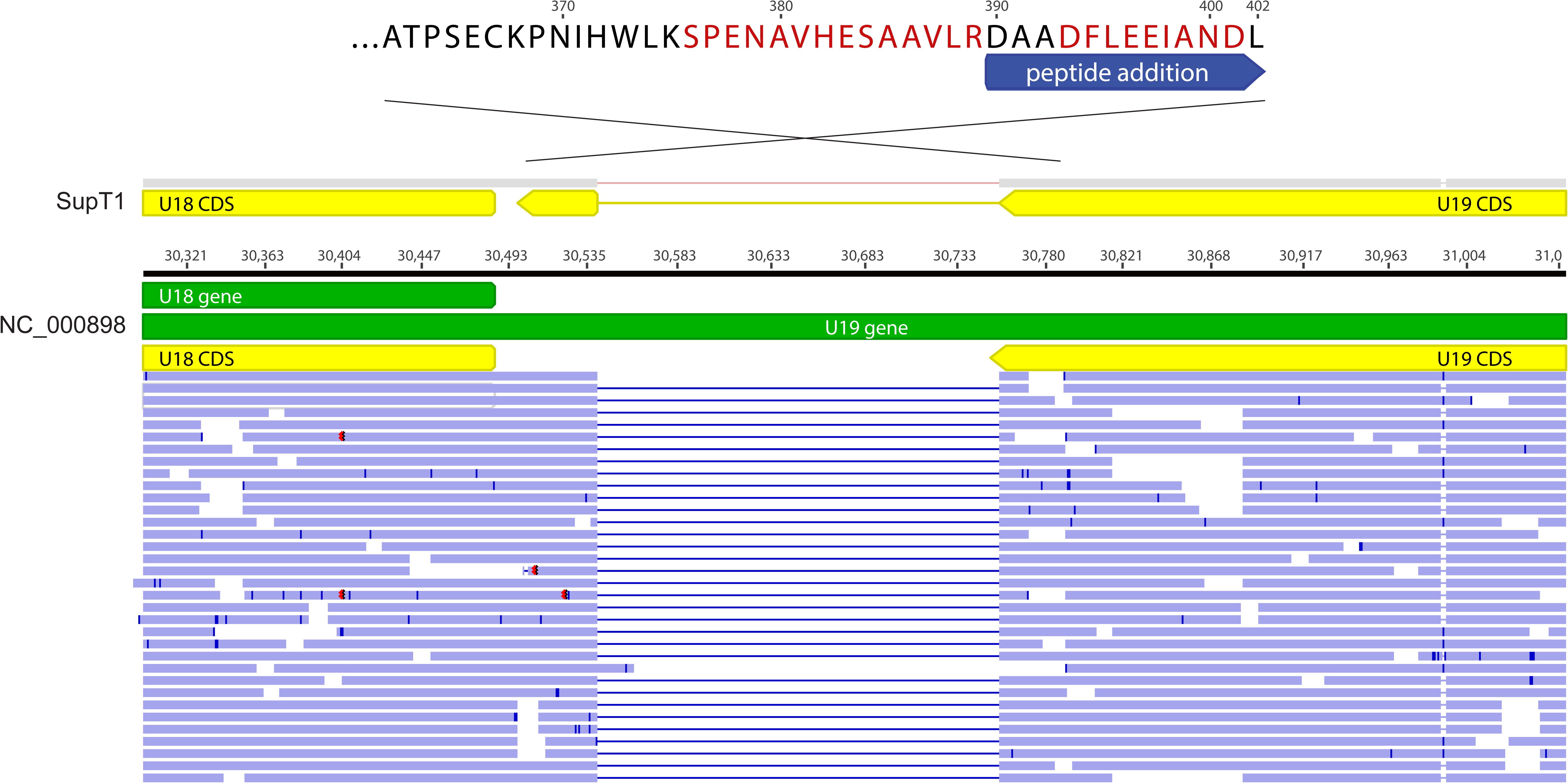
Unannotated splicing leading to C-terminal extension of HHV-6B U19 protein. Strand-specific RNA sequencing of HHV-6B cultured in Sup-T1 cells demonstrated a novel splice site at the 3’ end of the U19 transcript in the codon immediately before the annotated stop codon. The new splice site leads to a 13 amino acid C-terminal extension, which was confirmed by shotgun proteomics.

**Figure 9.**
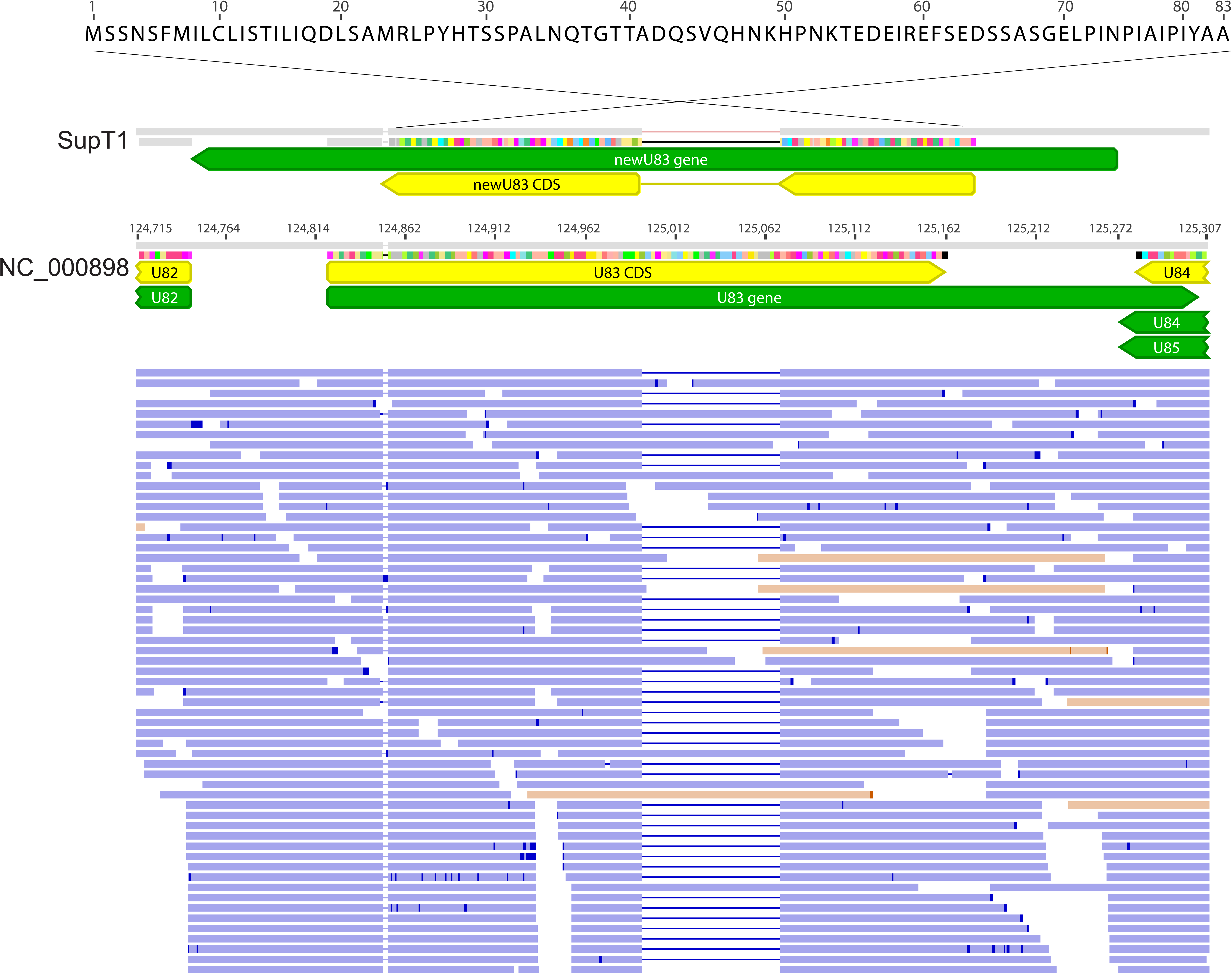
Antisense transcription and novel splicing of HHV-6B U83 gene. Nearly all of the strand-specific RNA-seq reads from Sup-T1 cells at the annotated HHV-6B Z29 U83 gene were antisense to the existing annotation and included a novel splice site. The same splice site in the context of antisense transcript predominance was recovered from virus cultured in MOLT3 cells. No high-confidence peptides were recovered to this alternatively spliced antisense transcript by shotgun proteomics.

## Discussion

In this study we sequenced 125 HHV-6B genomes and 10 partial HHV-6A genomes, increasing the full genome data available for HHV-6 by more than an order of magnitude. We found remarkably little sequence diversity among HHV-6B strains sampled from New York, Seattle, and Japan, with the average strain having fewer than 150 differences across the 119 kb unique long region relative to any other strain sequenced here. IciHHV-6B from across the United States had considerably less diversity than other cohorts of HHV-6 sampled. HHV-6A and HHV-6B strains sequenced here showed no overlap or recombination between species and the most divergent HHV-6B strain identified to date was isolated and sequenced. Viral sequences clustered by geographical origin and identical iciHHV-6B strains were found among many apparently unrelated individuals.

These results suggest that HHV-6B integration is a relatively infrequent event, that iciHHV-6B does not general reflect strains circulating in community causing acute infection, and that sequence diversity may be driven by a founder effect. Alternatively, certain strains could be prone to integration. At the same time, iciHHV-6B sequences were found admixed with HHV-6B strains from acute infection, suggesting that integration events are not uncommon. The hypothesis that HHV-6 integration into the germline is an infrequent event, however, would be consistent with a founder effect for each clade of identical iciHHV-6B found across our North American patients and account for the identical iciHHV-6 sequences found between two pairs of individuals from different sides of the Atlantic Ocean. It would also suggest that chromosomal integration of HHV-6 into the germ line is an extraordinarily rare event and almost all iciHHV-6 individuals acquired their virus from a remote integration event. More sequencing of both circulating HHV-6 strains and iciHHV-6 individuals is needed to test this hypothesis and will no doubt become available as more human genomes are sequenced. The hypothesis that integration bias due to viral sequence is the cause of the degeneracy of iciHHV-6 genomes is difficult to separate from founder effect and would only be testable in vitro or by following many individuals acutely infected with different strains of HHV-6B.

Despite widespread recombination, phylogenetic analyses demonstrated geographical clustering of HHV-6B strains with unique clades for Japanese strains and for several of the New York strains. Of note, the only patient of Asian descent in the New York cohort aligned best to the Japanese strains. These data would be consistent with the hypothesis of a familial source of transmission of acute HHV-6B. Because of the clustering of New York and Japan HHV-6 sequences, we are unable to ascertain whether strain differences can account for the striking differences in reported rates of encephalitis in infants with primary HHV-6 infection between Japan and the United States (30).

The geographical cluster of HHV-6B is similar to that seen for HSV-1 and HSV-2 genome sequences, which also show high degrees of interspecies recombination (25, 26, 31). The limited diversity of HHV-6B as measured by average pairwise nucleotide diversity is comparable to that found in HSV-2 in contrast to that identified in HSV-1 strains (25, 26). Of note, the diversity seen in HHV-6B is substantially less than that seen for the phylogenetically related human betaherpesvirus CMV (HHV-5) (27). No comparative genomics have been performed to date on the other human betaherpesvirus HHV-7.

Limitations of our approach include the limited worldwide sampling of HHV-6B strains, which included the Uganda, Japan, and the United States (with samples in the iciHHV-6 Fred Hutchinson cohort coming from several northern European individuals and only one Australian individual). Of note, our North American iciHHV-6 sequencing included individuals from at least 25 different states. More strains from both acutely infected and iciHHV-6 individuals are needed from Asia, the Middle East, South America, and Africa. Given the diversity seen in a limited subset of Ugandan strains and the limited diversity seen in iciHHV-6 in our study, it would be worthwhile to sequence iciHHV-6 from African populations to test hypotheses on the contribution of founder effect and strain sequence effects on HHV-6 integration. Sequencing of the U90 gene from reactivated HHV-6B strains from our clinical lab revealed additional lineages of HHV-6, which were subsequently confirmed by sequencing Ugandan HHV-6 isolates. Our clinical U90 sequences indicate even more lineages exist that we have not sampled on a genome-wide basis.

We also were not able to sequence through every repeat in the virus and thus our estimates of diversity would be biased to the null given that the repetitive elements may be one of the first sites of genome evolution. We also were not able to recover near-complete genomes of HHV-6A due to the use of a HHV-6B capture panel for sequencing. Future studies should be focused on continuing to probe the global diversity of HHV-6 sequences, understanding the degree of admixture between acute infections and iciHHV-6 strains, and whether genotypes identified here are associated with different clinical outcomes. Based on the results presented here, there was no clear association between viral sequence and clinical phenotypes such as CNS symptoms, although our power to detect such differences was limited.

Our RNA-sequencing data found novel spliceoforms and antisense transcripts in 10% of the genes currently annotated in HHV-6B Z29. Shotgun proteomic analysis recovered peptides for three changes in HHV-6B coding sequences and confirmed expression of 39 existing proteins in lytic HHV-6B infection. We also discovered differential splicing of U79 in SupT1 versus MOLT3 cells. These data allow for the most comprehensive annotation of an HHV-6 genome to date. Certainly, more work is also required to characterize how the novel spliceoforms, extensions, and transcripts discovered here affect viral replication and gene function, and whether they are present in the many strains sequenced here.

The sequences recovered here represent by far the largest HHV-6 sequencing effort conducted to date and increases the number of available genomes for HHV-6B by almost 60-fold. These data provide a wealth of information for clinical diagnostics, vaccine and antiviral design, and viral-host genome evolution. An immediate outcome of this resource has been to assist the development of a HHV-6B ORFeome project that will enable downstream studies in gene function and T-cell epitope and antigen discovery.

## Materials and Methods

### Collection of specimens

*New York cohort*

35 HHV-6B viral isolates were obtained from peripheral blood samples from children under 3 years of age with acute febrile illnesses or seizures presenting to the University of Rochester Medical Center Emergency Department or ambulatory settings in Rochester, NY, as previously described (4, 18, 32, 33). Samples from children with a known abnormality of immune function were excluded. Use of these samples was approved by the University of Rochester Institutional Review Board.

Peripheral blood mononuclear cells (PBMCs) were separated from EDTA anticoagulated blood samples via density gradient centrifugation (Histopaque 1077;Sigma Diagnostics, St. Louis, Mo.), and co-cultivated with stimulated cord blood mononuclear cells. Positive cultures were identified by characteristic cytopathic effect (CPE), confirmed by indirect immunofluorescent staining with monoclonal antibodies directed against HHV-6A and HHV-6B, and polymerase chain reaction, as previously described (4, 34).

### Japanese cohort

HHV-6B was isolated from PBMCs obtained from 10 ES patients and 10 HSCT recipients by co-cultivation with stimulated cord blood mononuclear cells. Infected cultures were identified on the basis of cytopathic effect (i.e., characteristics of pleomorphic, balloon-like large cells). The presence of virus was confirmed by immunofluorescence staining of the co-cultures with a specific HHV-6B monoclonal antibody (OHV-3; provided by T. Okuno, Department of Microbiology, Hyogo College of Medicine, Hyogo, Japan). Co-cultivated cord blood mononuclear cells infected with the clinical isolates were stored after several passages at −80 °C until assayed.

### Uganda cohort

Saliva samples were obtained from infants in a previously described birth cohort study of primary herpesvirus infection (35). Acute HHV-6B infection determined by weekly PCR testing of oral swabs. Whole saliva was collected every 4 months using the Salivette^®^ collection system (Sarstedt), transferred to cryovials, and frozen at −80°C until assayed. The samples used for this study were from 2 infants (both 3 months old at the time of sampling), 3 older children (ages 2.1 years, 2.8 years, and 4.2 years), and 1 adult (age unknown).

### IciHHV-6 cohort

74 individuals with iciHHV-6A or -6B were identified as part of a continuation of a previously described study (36). DNA was extracted from beta lymphoblastoid cell lines (LCLs) generated from Epstein-Barr virus infected peripheral blood mononuclear cells (PBMCs) obtained from hematopoietic cell transplant recipients and donors. Patients received HSCTs at Fred Hutchinson Cancer Research Center (FHCRC) in Seattle, WA. Donors were sourced from patient relatives and international bone marrow donor registries. We then used a pooling testing strategy as previously described using quantitative PCR (37) and droplet digital PCR (38) to identify individuals with iciHHV-6. A conserved region of the U94 gene was amplified to distinguish between species HHV-6A and HHV-6B.

### University of Washington Virology patient cohort

Samples from 21 different individuals previously found to be HHV-6 PCR positive were randomly selected from plasma submitted for testing in the Clinical Molecular Virology Laboratory at the University of Washington in 2014-2015. The majority of samples were from post-transplant-associated testing for suspected HHV-6 systemic infections. Eleven samples were from children <16 years of age (3-16 years old), and 10 were from adults (17-51 years old). Samples were from 13 males and 8 females. Five samples had viral loads <1,000 copies/mL (910, 740, 720, 550, and 480) while the remaining viral loads ranged from 1,000 – 53,000 c/mL. Of these 11 gave sufficient sequence after nested PCR to be included in downstream analyses. Use of these samples was approved by the University of Washington Institutional Review Board.

### DNA extraction and quantitative PCR and U90 sequencing

Approximately 5 µg of DNA were extracted from B-LCLs with iciHHV-6 and aliquoted at concentrations of ∼200 ηg/µL. DNA from the Japan and New York strains was extracted from 200 ul of viral culture using QIAamp 96 DNA kit (Qiagen) and eluted into 100 ul of AE buffer (Qiagen). To quantify the amount of HHV-6 and human DNA, 10ul of purified DNA was used to perform real-time quantitative PCR as described previously (36). Plasma samples from the University of Washington patient cohort were extracted using a MagnaPure LC (Roche) and MagnaPure LC DNA Isolate Kit with a starting volume of 200uL and elution volume of 100uL. The U90 locus was amplified using a nested PCR protocol as described previously (39). Amplicons from the same patient were pooled, diluted, and next-generation sequencing libraries were created using the Nextera XT kit.

### Sequencing of U91 RNA transcript

Seven million HHV6B (Z29)-infected SupT1 cells (from NIH AIDS Reagent Program) were used as starting material to create an RNA library with the Qiagen RNeasy Mini Kit according to manufacturer’s instructions. Total RNA was treated with TURBO DNase I (Thermo Fisher Scientific) and then used to create a cDNA library with SuperScript II Reverse Transcriptase (Thermo Fisher Scientific) according to manufacturer’s instructions. Using this cDNA as template and Platinum *Taq* DNA Polymerase High Fidelity (Thermo Fisher Scientific), the *U91* transcript was amplified by PCR with annealing temperature of 55.5° C for 30 cycles with primers that included cloning recognition sequences as follows: *U91* sense, 5’-GGGGACAAGTTTGTACAAAAAAGCAGGCTTCTCTGTAACACTGATCATGATGGGATATGAGGA-3’; *U91* antisense, 5’-GGGGACCACTTTGTACAAGAAAGCTGGGTCTTACACATTCATTTCAGTTTTC GGTATAATAGCCTC-3’. This PCR product was inserted into the pDONR221 cloning vector (Thermo Fisher Scientific) and Sanger sequenced using the M13F(-21) and M13R primers.

### Capture sequencing

Sequencing libraries for the New York, Japan, and iciHHV-6 cohorts were prepared using 100ng of genomic DNA using either NEB fragmentase, end repair/dA tailing, Y-adapter ligation, and dual-index Truseq PCR based or via the Kapa HyperPlus kit, following manufacturer’s protocol (40). Approximately 60ng of cleaned, amplified DNA library was pooled into sets of seven or eight samples based on relative viral qPCR to human beta-globin qPCR ratio, so that samples with similar relative concentrations of virus were pooled together. Capture sequencing was performed following the IDT xGen protocol with the use of half the amount of blocking adapter and at least 4 hours of 65C hybridization with a tiling biotinylated oligo capture library based on the reference HHV6-B genome (NC_000898). Post-capture libraries were sequenced to achieve at least 200,000 reads per sample library (at least 100X coverage based on at least 50% on-target) on a 1x180bp single-end run or on a 300x300bp paired-end run on an Illumina MiSeq

Capture sequencing for the Uganda cohort (n=6 samples) was performed using a custom-designed SureSelect^XT^ oligonucleotide panel covering HHV-6 and HHV-7 genomes and sequenced using an Illumina NextSeq using a v2 300 cycle mid-output kit (2x150bp paired end) (41, 42). Libraries were prepared as outlined in the SureSelect^XT^ Automated Target Enrichment protocol version J0 (December 2016) with two minor modifications. 20ng of total DNA was sheared prior to end-repair, A-tailing and adapter ligation (1:100 dilution). Two extra cycles of PCR were performed during library amplification prior to hybridization while four extra cycles of PCR were added to the post-hybridization amplification / indexing step.

### RNA-Seq of HHV-6B Z29 strain

Total RNA was extracted from MOLT3 and Sup-T1 cells asynchronously infected with HHV-6B Z29 strain with >10^6^ copies/mL of virus in the supernatant. 3ug of total RNA was used as input for polyA-purification and strand-specific RNA-Seq libraries were prepared from using the NEBNext Ultra Directional RNA Library Prep Kit. Two libraries were prepared from infected SupT1 cells and one from infected MOLT3 cells. Transcriptome libraries were sequenced on an Illumina MiSeq using multiple runs types (2x94bp, 1x188bp).

### Shotgun proteomics

Proteomic samples were prepared from soluble cell lysates or serum-free conditioned media from HHV6-infected Sup-T1 cells. HHV-6B quantitation in lysates was 23,683,766 copies per PCR reaction with a corresponding beta-globin copy number of 12,900 copies per reaction; HHV-6B quantitation in the serum-free media was 3,122,307 copies per reaction with a corresponding beta-globin copy number of 10,115 copies per reaction. Approximately 2-20 micrograms of protein were separated on two 10–20% Criterion Tris-HCl run in MOPS or one 4-12% Criterion Tris-HCl run in MES SDS-PAGE gels (Bio-Rad), silver stained, and gel bands were excised for mass spectrometry-based peptide sequencing as described previously (43) (S3 Figure). Samples were digested with sequencing grade trypsin (Promega) only, or with trypsin followed by AspN (Roche) following the standard UCSF MS facility protocol (http://msf.ucsf.edu/protocols.html) (44).

Peptide sequencing was performed using an LTQ-Orbitrap Velos (Thermo) mass spectrometer, equipped with a 10,000 psi nanoACUITY (Waters) UPLC. Reversed phase liquid chromatography was performed using an EasySpray C18 column (Thermo, ES800, PepMap, 3 µm bead size, 75 µm x 15 cm). The LC was operated at 600 nL/min flow rate for loading and 300 nL/min for peptide separation over a linear gradient over 60 min from 2% to 30% acetonitrile in 0.1% formic acid. For MS/MS analysis on the LTQ Orbitrap Velos, survey scans were recorded over 350-1400 m/z range, and MS/MS HCD scans were performed on the six most intense precursor ions, with a minimum of 2,000 counts. For HCD scans, isolation width was 3.0 amu, with 30% normalized collision energy. Internal recalibration to a polydimethylcyclosiloxane (PCM) ion with m/z = 445.120025 was used for both MS and MS/MS scans (45).

Mass spectrometry centroid peak lists were generated using in-house software called PAVA, and data were searched using Protein Prospector software v. 5.19.1 (46). Data were searched with carbamidomethylation of Cys as a fixed modification, and as variable modifications, oxidation of methionine, N-terminal pyroglutamate from glutamine, start methionine processing, and protein N-terminal acetylation. Trypsin, or trypsin plus AspN specificity was chosen as appropriate for each experiment. Mass accuracy tolerance was set to 20 ppm for parent and 30 ppm for fragment masses. For protein identification, searches were performed against a 9,874 entry database containing all protein sequences longer than or equal to 8 amino acids derived from HHV-6 Z29 strain genomic sequence translated in all six reading frames combined with translated splice junctions derived from RNA-Seq data. Searches were also performed with the SwissProt human database (*downloaded September 6, 2016*) containing 20,198 entries, and fetal bovine serum (P02769) as a cell culture supplement. Databases were concatenated with matched, fully randomized versions of each database to estimate false discovery rate (FDR) (47).

The HHV-6B protein database was searched initially allowing for two missed and one non-specific cleavage to allow for peptides with alternative splicing or unpredicted start/stop sites. Standard Protein Prospector scores (minimum protein score 22, minimum peptide score 15, maximum protein expectation value 0.01 and maximum peptide expectation value 0.001) produced a 5% FDR for protein identifications. All matched HHV6 peptide spectra were manually *de novo* sequenced, and may be viewed with the freely available software MS-Viewer, accessible through the Protein Prospector suite of software at the following URL: http://prospector2.ucsf.edu/prospector/cgi-bin/msform.cgi?form=msviewer, with the search key: 7awn6ehwzd. Raw mass spectrometry data files and peak list files have been deposited at ProteoSAFE (http://massive.ucsd.edu) with accession number MSV000081332.

### Sequence analysis

DNA Sequencing reads were quality and adapter-trimmed using Trimmomatic v0.36 and Cutadapt, *de novo* assembled using SPAdes v3.7 and mapped to reference genomes NC_000898 and NC_001664 using Bowtie2 (48– 50). Contigs were aligned to reference genomes using the multiple alignment program Mugsy v1.2.3 and resolved against consensus sequences from mapped reads using custom scripts in R/Bioconductor (51–53). Final assemblies were generated after discarding any contigs with mapq <= 5. Assembled genomes were annotated using Prokka and deposited to Genbank (accession numbers in S1 Table).

As the sequencing length was not sufficient to regularly discern sequence in the direct repeats and across several of the smaller repeats present in the HHV-6B genome, analysis was performed on aligned sequences that were pruned to keep four non-repeat-containing regions: between R0 and R1 repeats (UL), between R1 and R2A repeats (upstream and N-terminal U86 region), between R2B and R3 repeats (containing U90/91 genes), and between U94-U100 genes (Figure 1). Population genomics analyses including nucleotide diversity estimates, Tajima’s D, Achaz’s Y, and Hudson-Kaplan recombination estimates were executed using the PopGenome R package (28, 29, 54). Recombination detection analyses were performed using the DualBrothers package using a window length of 800 bp and a step size of 100 bp (55).

RNA sequencing reads were trimmed using cutadapt and mapped to the HHV-6B Z29 reference genome using Geneious v9.1 read aligner with structural variant discovery (decreased gap penalty) (56). RPKM values were calculated based on HHV-6B Z29 reference genome annotations and displayed using custom scripts in R/Bioconductor.

## Acknowledgements

Mass spectrometry analysis was provided by the UCSF Mass Spectrometry Facility directed by Al Burlingame, supported by the Adelson Medical Research Foundation. We appreciate assistance from Alex Yamana, Gabby Dolgonos and Krithika Nathamuni for careful inspection of peptide mass spectral assignments. We thank Samia Naccache and Dharam Ablashi for helpful comments on the manuscript.

## Figure Legends

**S1 Figure – Resequencing of select iciHHV-6B specimens confirms identical sequences among unrelated patients.** Samples from select iciHHV-6B specimens with identical sequences were re-extracted, re-prepared and re-sequenced from original patient material to rule out contamination or a sample specimen switch during the sequencing process. 11/12 of specimens gave identical sequence throughout the unique long region directly from de novo assembly. One specimen (ciHHV-6B-30E3) had one nucleotide change (G77564T) upon resequencing at a base that had a G/T variant allele frequency of approximately 50% each time the sample was sequenced.

**S2 Figure – Phylogenetic tree of HHV-6B complete U90/91 and U94/100 loci.** HHV-6B genomes were aligned using MAFFT, curated for sequence outside of repeat regions, and phylogenetic trees were constructed using MrBayes along the 6kb U90/91 (A), and 10kb U94-100 (B) regions. HHV6-6B NY310 was used as an outgroup. Samples are colored and labeled for origin based on New York (green), Japan (blue), or iciHHV6-B from HSCT recipients or their donors in Seattle (black), as well as whether two genomes were recovered from first-degree relatives (red).

**S3 Figure** - Non-contiguous gel images of silver stain of HHV-6B Z29 lysates in SupT1 cells or serum-free supernatant run on 10-20% TrisHCl gels in MOPS buffer

**S4 Figure** - Gel image of silver stain of HHV-6B Z29 lysate in SupT1 cells or serum-free supernatant run on 4-12% TrisHCl gel in MES buffer

